# An ABA Synthesis Enzyme Allele *OsNCED2* Promotes the Aerobic Adaption in Upland Rice

**DOI:** 10.1101/2020.06.11.146092

**Authors:** Liyu Huang, Yachong Bao, Shiwen Qin, Min Ning, Jun Lyu, Shilai Zhang, Guangfu Huang, Jing Zhang, Wensheng Wang, Binying Fu, Fengyi Hu

## Abstract

There are two ecotypes, upland rice and irrigate rice, during its evolution in rice. Upland rice exhibits aerobic adaptive phenotype via its stronger root system and more rapid drought responses compared with its counterpart, irrigated rice. We assessed the functional variation and applications of the aerobic adaptive allele *OsNCED2^T^* cloned from IRAT104 which is a cultivar of upland rice. *OsNCED2*-overexpressing transgenic rice and *OsNCED2^T^*-NILs exhibited significantly higher ABA contents at the seedling and reproductive stages, which can improve root development (RD) and drought tolerance (DT) to promote aerobic adaptation in upland rice. RNA-Seq-based expression profiling of transgenic versus wild-type rice identified *OsNCED2-*mediated pathways that regulate RD and DT. Meanwhile, *OsNCED2*-overexpressing rice exhibited significantly increased reactive oxygen species (ROS)-scavenging abilities and transcription levels of many stress- and development-related genes, which regulate RD and DT. A SNP mutation (C to T) from irrigated rice to upland rice, caused the functional variation of *OsNCED2*, and the enhanced RD and DT mediated by this site under aerobic conditions could promote higher yield of upland rice. These results show that *OsNCED2^T^*, through ABA synthesis, positively modulates RD and DT which confers aerobic adaptation in upland rice and might serve as a novel gene for breeding aerobic adaptive or water-saving rice.

## INTRODUCTION

Rice is a major food crop and model plant for the genomic study of monocotyledon species. Long-term natural selection has driven the evolution of various ecotype populations in *Oryza sativa*, such as upland rice and irrigated rice, which have evolved their respective characteristic traits under both natural and artificial selection, which leads to phenotypic adaptation to the respective environment and simultaneous genetic differentiation between ecotypes. Therefore, rice provides a model system for studying adaptation to aerobic and drought environments (Xia et al., 2019). Although the yield of upland rice is comparatively lower than that of irrigated rice, the former serves as a daily staple food on which nearly 100 million people depend (Pomfret, 1986). Accordingly, upland varieties are widely utilized under upland conditions to optimize crop yields due to their more efficient usage of water and their adaptation to aerobic upland (Xia et al., 2019). Previous research has proposed three different mechanisms for plant drought resistance: drought escape, dehydration avoidance, and drought tolerance (DT) (Jérôme et al., 2010). Upland rice has multiple special characteristics, such as aerobic adaption, taller height, lower tillering potential, and longer and denser roots, compared with its irrigated counterpart (Chang, 1982; Chang and Vergara, 1975).

In our previous study, the genomes of upland and irrigated rice accessions were analysed via resequencing and population genetics (Lyu et al., 2014). Multiple pathways or ecotype differentiated genes (EDGs) that potentially contribute to the aerobic adaptation of upland rice to dryland were identified, but only a few of these have been functionally verified. Interestingly, a 9-*cis*-epoxycarotenoid dioxygenase gene (*Os12g0435200*) named *OsNCED2*, which plays key roles in the abscisic acid (ABA) synthesis pathway, was identified as a domestication gene which contributes to the aerobic adaptation of upland rice (Lyu et al., 2013). The ABA-mediated root tropic response and stomatal closure under water stress are key responses of plants that ensure their survival under water-limited conditions. ABA signalling also plays a critical role in the regulation of root growth and the architecture of the root system and likely interacts with other hormones, such as auxins, gibberellins, or brassinosteroids (BRs), to regulate in these processes (Deak and Malamy, 2005; Rodrigues et al., 2009; Swarup et al., 2005). ABA is a stress-responsive phytohormone that inhibits seed germination and seedling growth to adapt to unfavourable environmental conditions, whereas gibberellic acid (GA) and BRs are major growth-promoting phytohormones that promote seed germination, seedling growth, flowering and leaf expansion (Clouse, 2016; Cutler et al., 2010; Golldack et al., 2013; Wang et al., 2018; Weiner et al., 2010).

In this study, *OsNCED2* was functionally identified as a 9-*cis*-epoxycarotenoid dioxygenase that mediates ABA synthesis. Furthermore, *OsNCED2* can promote root development while reducing plant height in *OsNCED2*-overexpressing transgenic lines. Although both types of OsNCED2 (C/T) have a 9-*cis*-epoxycarotenoid dioxygenase function, they might exhibit significant differences in catalytic activity, which result in the different ABA concentrations between upland rice and irrigated rice. Thus, rice with T-type *OsNCED2*, *OsNCED2^T^*, exhibits increased adaptation to aerobic conditions due to its higher ability to regulate the synthesis and/or distribution of ABA in response to osmotic stress. Consequently, both the overexpression of *OsNCED2* and the import of *OsNCED2^T^* into irrigated rice enhance plant yield and therefore can potentially be used in the breeding of aerobic adaptive or water-saving rice.

## RESULTS

### Identification and Characterization of *OsNCED2* from Upland Rice and Irrigated Rice

In our previous comparative genomic study, we found that *OsNCED2* on chromosome 12, was highly differentiated between the two ecotypes (Lyu et al., 2013). *OsNCED2* differs in abundance between upland and irrigated rice and produces two haplotypes of the SNP (C/T), namely, T-type in upland rice and C-type in irrigated rice (Lyu et al., 2013). The full-length *OsNCED2* gene sequence consists of 1731 bp without any intron and encodes a 576-amino-acid polypeptide annotated as a 9-*cis*-epoxycarotenoid dioxygenase. A phylogenetic analysis indicated that all NCED paralogous genes might have originated from two ancestral genes in rice and that *OsNCED2* clustered with other paralogous genes, which implies that *OsNCED2* might have arisen from gene duplication in the rice genome (Supplemental Fig. S1). *OsNCED2* contains several signal-responsive *cis*-elements in its promoter region (1.5 kb upstream of the start codon), and these include light, anaerobic, GA, MeJA and ABA response elements (Supplemental Table S1), which implies that the expression of *OsNCED2* could be regulated by environmental and hormone signalling. A Target P analysis confirmed that OsNCED2 might be located in chloroplasts and our cytological analyses indicate that OsNCED2 is localized in the chloroplast (Supplemental Fig. S2), which is consistent with its function as a 9-*cis*-epoxycarotenoid dioxygenase that catalyses the synthesis of ABA in chloroplasts (Qin and Zeevaart, 1999).

### Spatial-temporal Expression Pattern of *OsNCED2* in Rice

Because *OsNCED2* has seven paralogous genes in the whole genome and might play different roles in rice development and response to the environment, we examined the *OsNCED2* expression patterns in different tissues at different developmental stages by both qRT-PCR and histochemical staining, which indicated the occurrence of *OsNCED2* promoter activity in *OsNCED2Pro::GUS* reporter transgenic plants. The results showed that *OsNCED2* was mainly expressed in the mature leaves, stems and roots and exhibited relatively lower expression in young panicles and seedlings (Fig. 1, A). Although differences in the *OsNCED2* promoter were found between upland and irrigated rice, no significant differences in expression was detected between the upland rice variety IRAT104 and the irrigated rice variety Yueguang (Fig. 1, B). GUS histochemical staining of *OsNCED2Pro::GUS* reporter transgenic plants also showed consistent expression in related tissues (Fig. 1, C). Interestingly, *OsNCED2* was mainly expressed in new or young but not old roots (Fig. 1, D). These results suggest that *OsNCED2* might play different roles in different tissues.

**Figure 1.**
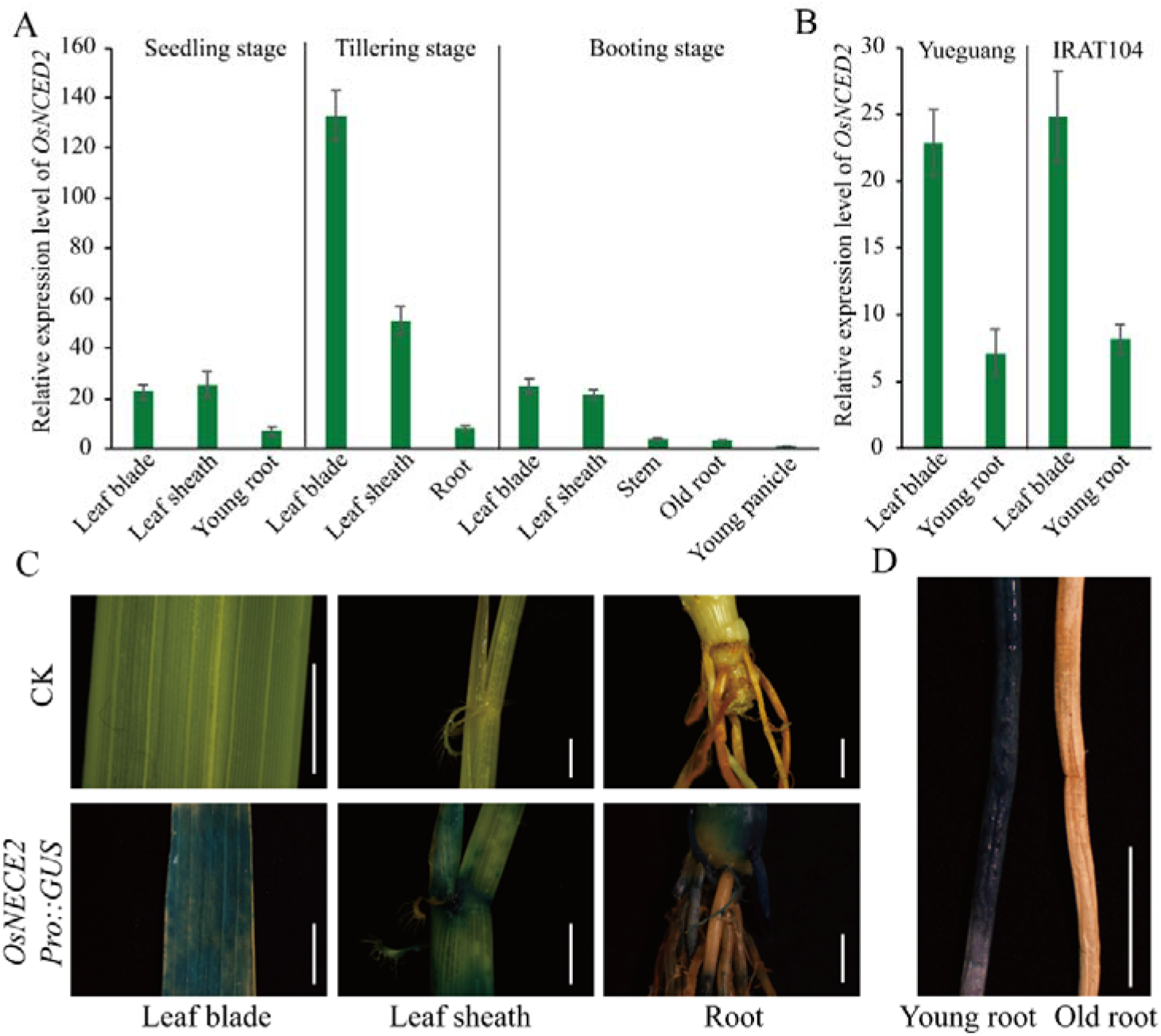
Expression pattern of *OsNCED2* in different tissues of rice. A Relative expression level (2^−ΔΔCt^) of *OsNCED2* in different rice tissues at the seedling, tillering and booting stages. Young panicles were used as reference controls. B Koshihikari and IRAT104 exhibit similar *OsNCED2* expression levels and models. C *OsNCED2* expression in different tissues of *OsNCED2Pro::GUS* transgenic rice plants determined by GUS staining analysis. D Novel young roots exhibit high *OsNCED2* expression (left), whereas old roots show low *OsNCED2* expression (right). Scale bars, 0.5 cm.

### *OsNCED2* Overexpression can Promote ABA Synthesis and Root Development

To verify the function of *OsNCED2*, two types of *OsNCED2* alleles cloned from IRAT104 (upland rice with *OsNCED2^T^*) and Nipponbare (irrigated rice with *OsNCED2^C^*) were overexpressed in Nipponbare (Fig. S3). An analysis of the root phenotype showed that the overexpression of both *OsNCED2^T^* and *OsNCED2^C^* can promote crown root development, which results in a denser root system at both the seedling and tillering stages (Fig. 2, A and B; Table 1). Moreover, the overexpression of both *OsNCED2^T^* and *OsNCED2^C^* can enhance the ABA level compared with that found in non-transgenic plants (Fig. 2, C and D). The lack of a positive correlation between the expression of *OsNCED2^T^* or *OsNCED2^C^* (Fig. 2, E) and the ABA content in the transgenic plants implies that the catalytic activity of OsNCED2^T^ is higher than that of OsNCED2^C^, which is consistent with the result that near-isogenic lines (RILs) with *OsNCED2^T^* exhibit significantly higher ABA levels and denser lateral roots than C-type RIL families (Lyu et al., 2013). Plant root injury discharge can indicate the vitality and water-absorption ability of roots. Therefore, we detected the plant root injury discharge of the transgenic lines overexpressing *OsNCED2^T^* or *OsNCED2^C^*. As shown in Fig. S4, almost all the transgenic lines exhibited significantly higher injury discharge than the WT plants. These results further prove that *OsNCED2* can mediate rice root development by catalysing ABA synthesis.

**Figure 2.**
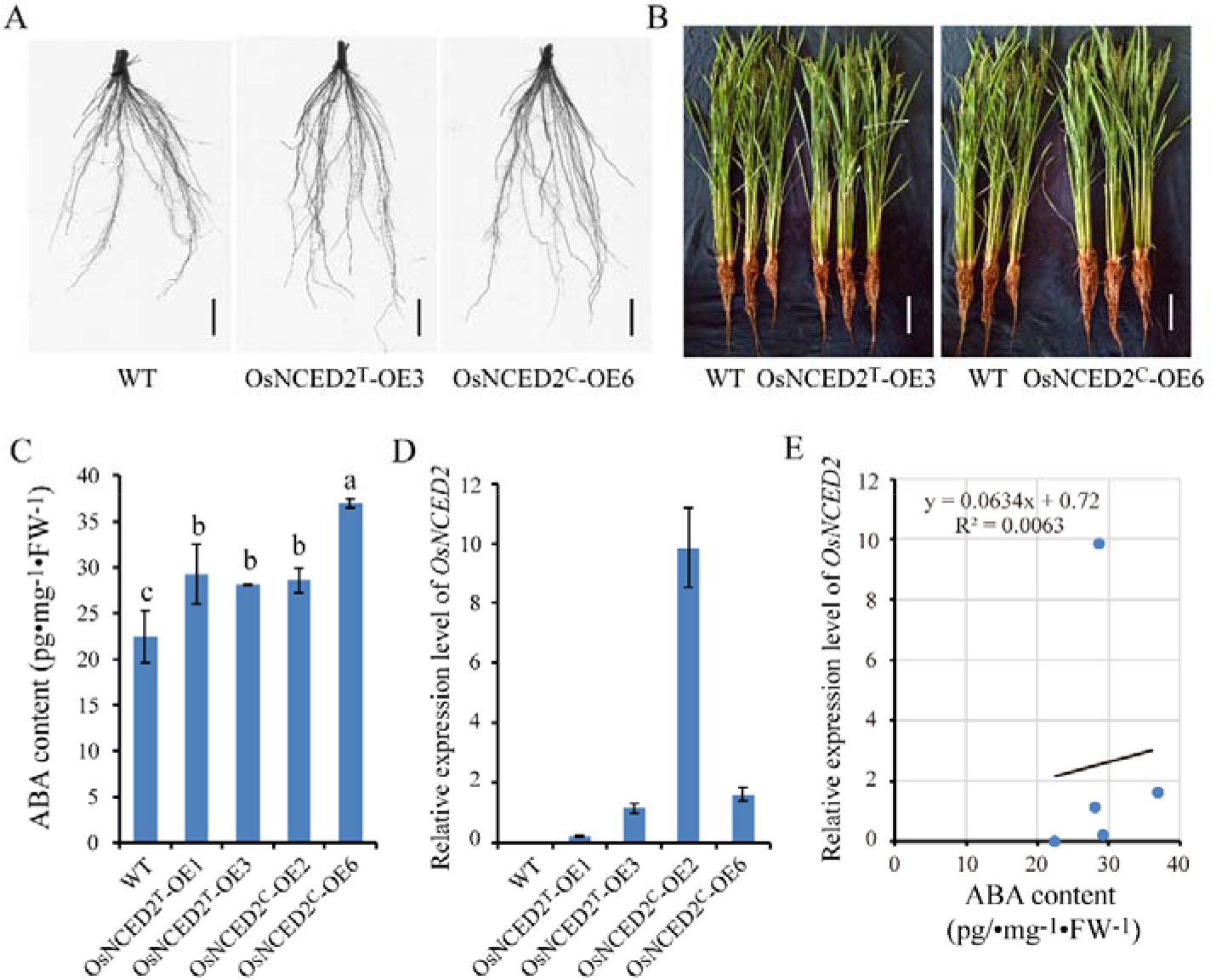
Both *OsNCED2^T^* and *OsNCED2^C^* can promote ABA synthesis and enhance root development. Root phenotype (A) and plant phenotype (B) of transgenic rice overexpressing both *OsNCED2^T^* and *OsNCED2^C^*. C ABA content in leaves of *OsNCED2^T^*-OE and *OsNCED2^C^*-OE transgenic plants. D Expression level (2^−ΔCt^) of *OsNCED2* in transgenic rice. The ABA content exhibited no correlation with the *OsNCED2* expression levels in lines overexpressing *OsNCED2^T^* and *OsNCED2^C^*. Scale bars, 2 cm (A), 20 cm (B).

### *OsNCED2* Overexpression can Enhance the Drought Tolerance of Rice

Stomatal closure is one of the first responses to drought conditions that might control plant dehydration. The stomatal status of the *OsNCED2^T^*-OE1 and OE3 leaf surfaces at the seedling stage under both normal and drought stress was examined by scanning electron microscopy (SEM), and the results indicated that the leaf surfaces of the *OsNCED2^T^*-OE1 and OE3 plants exhibited significantly higher stomatal closure rates than those of the WT plants (Fig. 3, A).

**Figure 3.**
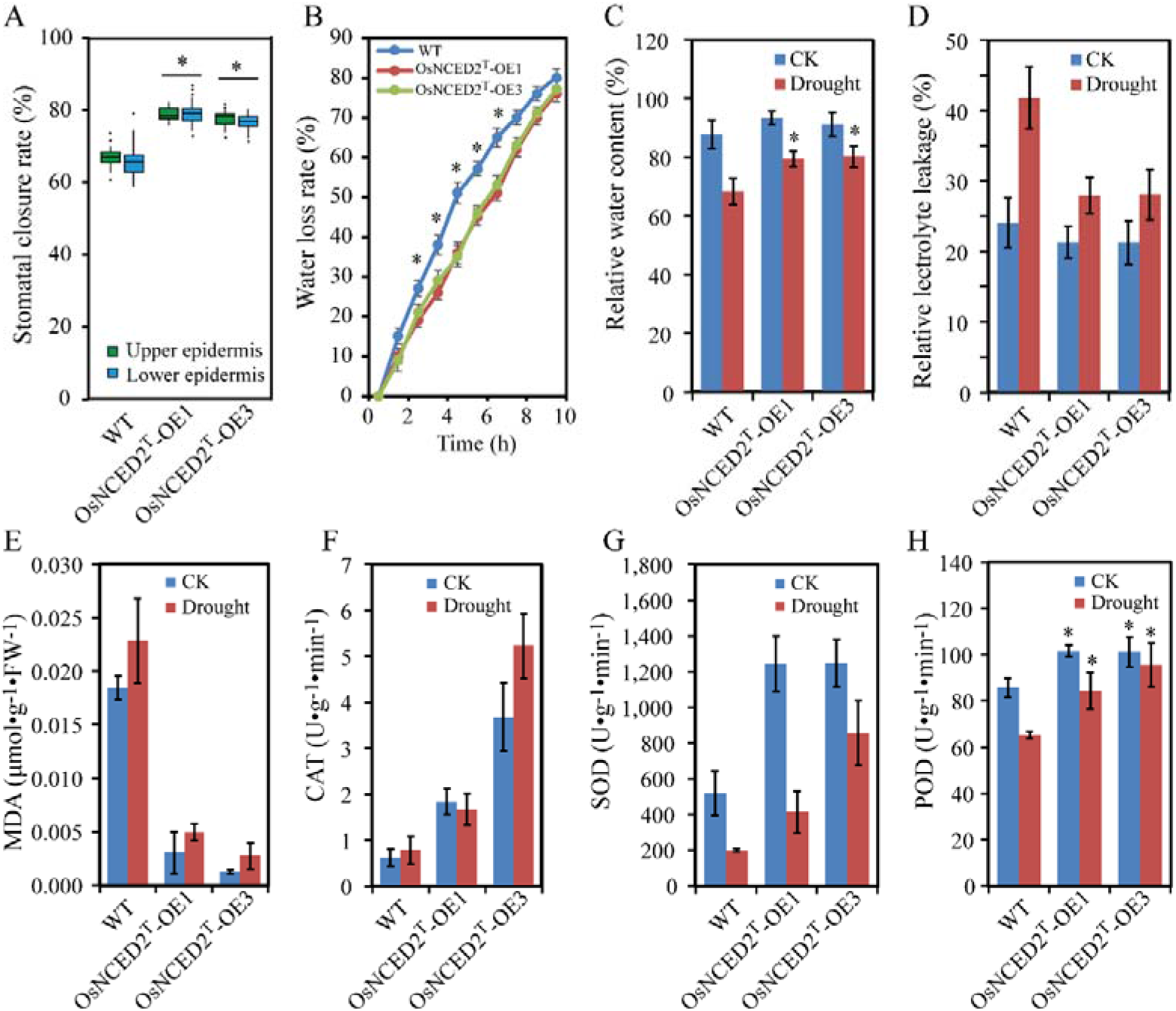
Physiological traits of *OsNCED2^T^*-OE lines under normal (CK) conditions and 1-h drought treatment. A Stomatal closure rate of leaves of *OsNCED2^T^*-OE and WT lines (both the upper and lower epidermis were counted in 10 randomly selected high-power fields). B Water loss rate of the transgenic and WT lines. From each replicate, 30 fully expanded leaves at the seedling stage were used in a triplicate experiment. Relative water content (C) and relative electrolyte leakage (D) of *OsNCED2^T^*-OE lines under normal conditions and drought treatment. (E) MDA contents of *OsNCED2^T^*-OE and WT plants under normal conditions and after 1 h of drought treatment. Activity of ROS-scavenging enzymes, including CAT (F), SOD (G) and POD (H), in *OsNCED2^T^*-OE and WT plants under normal conditions and after 1 h of drought treatment. The values are presented as the means and standard errors from three replicates. \**p* < 0.05 versus the WT plants, as determined by Student’s *t*-test.

To investigate the physiological responses mediated by *OsNCED2^T^* and *OsNCED2^C^*, several indices of drought-induced effects on OE and WT leaves at the shooting stage under both normal and drought conditions were measured. The water loss rate (WLR) from excised leaves was determined, and the results showed that the OE leaves showed a relatively lower WLR than the WT leaves (Fig. 3, B). Accordingly, the relative water contents (RWCs) in the *OsNCED2^T^*-OE1 and OE3 plants after 1 h of exposure to drought stress (77.9% and 78.6%) were significantly higher than that in the WT plants (67.4%) (Fig. 3, C). The OE seedlings experienced less extensive cell membrane injury (relative electrolyte leakage) and exhibited lower malondialdehyde (MDA) concentrations after drought treatment compared with the WT seedlings (Fig. 3, D and E). Additionally, under drought stress conditions, the catalase (CAT), POD and SOD activities in the *OsNCED2^T^*-OE1 and OE3 plants were significantly higher than those in the WT plants (Fig. 3, F, G and H), which indicates that the *OsNCED2^T^*-OE plants exhibit active detoxification by reactive oxygen scavenging regulation in response to drought (Sofo et al., 2015). Taken together, these results demonstrate that the DT of *OsNCED2^T^*-OE1 and OE3 plants was significantly improved when compared with that of the WT plants, which indicates that the overexpression of *OsNCED2* can enhance DT via physiological regulation.

### *OsNCED2^T^*-NILs Exhibit Higher ABA Levels and More Developed Root Systems than the *OsNCED2^C^* Background

Although RILs with *OsNCED2^T^* display a significantly higher ABA content and denser lateral roots than C-type RIL families (Lyu et al., 2013), other loci can influence the phenotype. To evaluate whether only the *OsNCED2^T^* locus contributes to ABA enhancement and root development, we generated three NILs named T-Koshihikari-1~3 that contain the *OsNCED2^T^* locus in the Koshihikari genetic background (C-Koshihikari), which was selected from the BC_4_F_9_ population lines by marker assisted selection (MAS) with a CAPs marker (Supplemental Table S2). A root phenotype analysis indicated that NILs with *OsNCED2^T^* contained a stronger root system, particularly during crown root development (Fig. 4, A; Table 2). Interestingly, although the leaf ABA content was markedly higher than that in the root, the ABA level showed sharp increases in the root but not the leaves in both genotypes in response to drought stress (Fig. 4, B). Furthermore, the ABA content was significantly higher in the T-Koshihikari-1 leaves than in the C-Koshihikari leaves under both normal and drought conditions (Fig. 4, B). In contrast, the ABA content in the T-Koshihikari1 roots was significantly higher than that in the C-Koshihikari roots only under drought conditions (Fig. 4, B). To clarify the DT of the *OsNCED2^T^*-NIL plants, we compared the growth status and survival rate of Koshihikari and *OsNCED2^T^*-NIL seedlings under polyethylene glycol (PEG) treatment. When the *OsNCED2^T^*-NIL and C-Koshihikari seedlings were grown in hydroponic culture under osmotic stress imposed by PEG, the *OsNCED2^T^*-NIL plants also exhibited improved DT compared with the C-Koshihikari plants (Fig. 4, C and D), which suggests that OsNCED2^T^ contributes to dehydration tolerance in plants. These results indicate that *OsNCED2^T^* might promote root development by regulating ABA synthesis under drought conditions and that this effect might contribute to drought avoidance and DT, which is as a kind of aerobic adaptation in upland rice.

**Figure 4.**
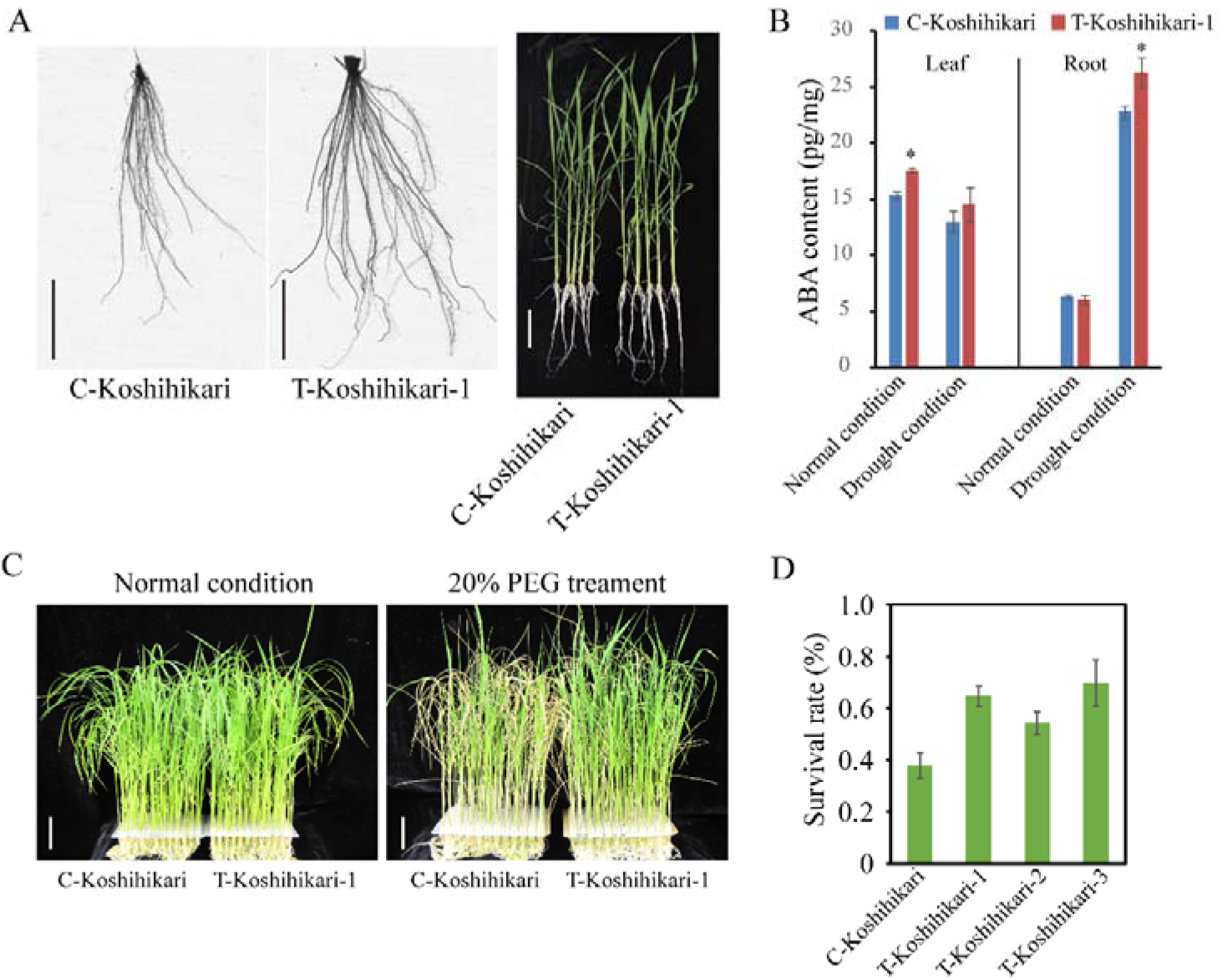
*OsNCED2^T^*-NIL plants have an enhanced root system and a stronger ABA synthesis ability compared with the background C-Koshihikari plants with *OsNCED2^C^*. A Plant and root phenotypes of *OsNCED2^T^*-NIL and C-Koshihikari plants. B Leaf and root ABA contents in *OsNCED2^T^*-NIL and C-Koshihikari plants under both normal conditions and drought treatment. C Phenotype of *OsNCED2^T^*-NIL and C-Koshihikari plants under both normal and drought (1 week of treatment with 20% PEG treatment) conditions. D Survival rate under re-watered conditions after 1 week of PEG stress treatment. Scale bars, 5 cm.

### Under Aerobic Conditions with Drought Stress, *OsNCED2^T^* can Enhance Drought Tolerance, which Results in Higher Yield

To investigate the effect of *OsNCED2^T^* and its potential application in rice breeding, we compared the grain yields of C-Koshihikari and T-Koshihikari plants under normal and aerobic conditions with no, moderate (natural upland condition) and severe drought (artificially controlled by rain proof installations). Under normal conditions, both lines exhibited similar yields, and no significant differences in other agronomic traits except the panicle number per plant were detected (Fig. 5, A, B, C and D; Supplemental Table S3). Although moderate drought significantly reduced the tiller number (even panicle per plant), percentage of filled grains and 1000-grain weight in all the tested plants, the *OsNCED2^T^*-NIL plants exhibited a higher grain weight per plant than the C-Koshihikari plants because of the above three yield-related factors (Fig. 5, A, B, C and D). Under severe drought, more prominent physiological damage, such as leaf wilting and delayed flowering, was detected in the C-Koshihikari plants compared with the *OsNCED2^T^*-NIL plants (Fig. 5, G; Supplemental Table S3). Furthermore, the yield per plant or unit area of the *OsNCED2^T^*-NILs was significantly higher than that of the C-Koshihikari plants (Fig. 5; Supplemental Fig. S5). These results confirm that under drought conditions, both the enhanced rooting and the dehydration tolerance promoted by *OsNCED2^T^* facilitate improvements in growth vigour and grain filling, which results in higher yield.

**Figure 5.**
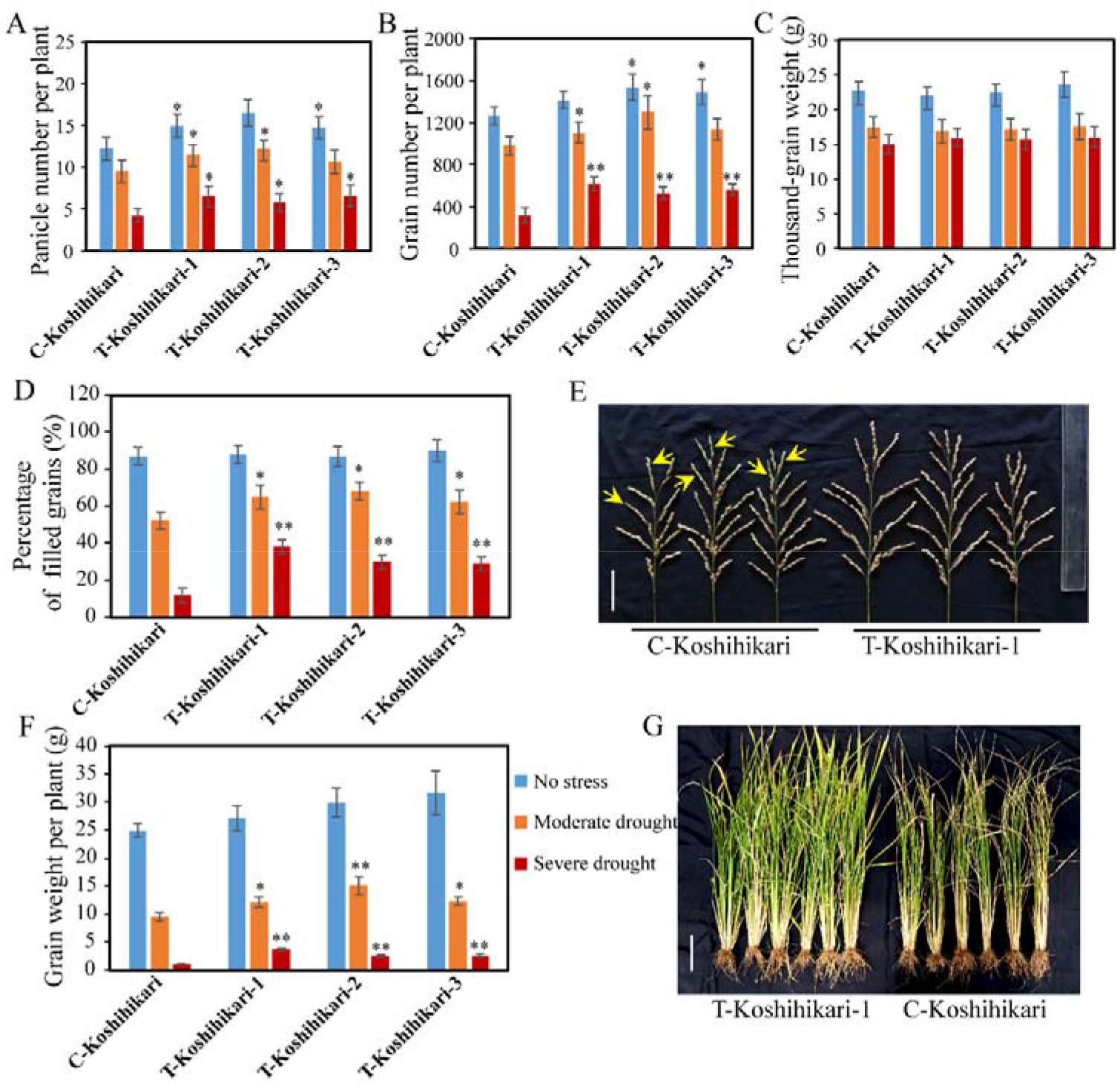
Effect of *OsNCED2^T^* on yield during exposure to aerobic and drought stress. Panicle number per plant (A), grain number per plant (B), thousand-grain weight (C), percentage of filled grains (D) and grain weight per plant (F) obtained for C-Koshihikari and *OsNCED2^T^*-NIL plants grown during exposure to no, moderate and severe drought. The values in the histograms represent the means ± SDs (n = 24). The panicle number refers to the number of effective tillers that were fertile. The panicle phenotypes (E) include the primary and secondary branch numbers and the grain number under moderate-drought conditions and the plant performance (G) under severe-drought conditions. The yellow arrows indicate abortive grains. * and ** indicate significant differences from the C-Koshihikari plants at *p* < 0.05 and *p* < 0.01, respectively, as determined by Student’s *t*-test. Scale bars, 5 cm (E), 10 cm (G)

### The SNP Mutation C to T Contributes to the Functional Differentiation of *OsNCED2* between Irrigated Rice and Upland Rice

To define whether the SNP mutation, which is C-to-T from irrigated rice to upland rice is responsible for the functional variation of *OsNCED2*, a further mutation analysis was conducted with the aim of verifying whether the SNP that produces the amino acid change from valine to isoleucine results in the observed difference in catalytic activity and plays important roles in the ecotypic differentiation of upland rice. A newly derived CRISPR/Cas9 gene editing technology was used to edit the C-SNP of *OsNCED2* in Nipponbare (see the Materials and Methods section). Eight positive independent lines identified as heterozygous in this site were identified from 32 transgenic T_0_ lines via both the CAPs marker and Sanger sequencing. The root system and DT of both T-type and C-type Nipponbare plants belonging to the T_1_ generation were evaluated. The results indicated that the T-type plants had a more enhanced root system than the C-type plants at both the seedling (Fig. 6, A) and mature stages (Fig. 6, B). Furthermore, the yield per plant showed no significant difference between the T-type and C-type Nipponbare plants under normal conditions (Fig. 6, C). Consequently, under upland conditions, the final grain weight per plant obtained for the T-type plants was significantly higher than that of the C-type plants (Fig. 6, C). These results confirm that the SNP mutation site C-to-T from irrigated rice to upland rice results in the functional variation of *OsNCED2*, and that both the enhanced rooting and the dehydration tolerance mediated by this site promote a higher yield under drought conditions, which indicates that this site plays vital roles in the adaptation and artificial selection of upland rice.

**Figure 6.**
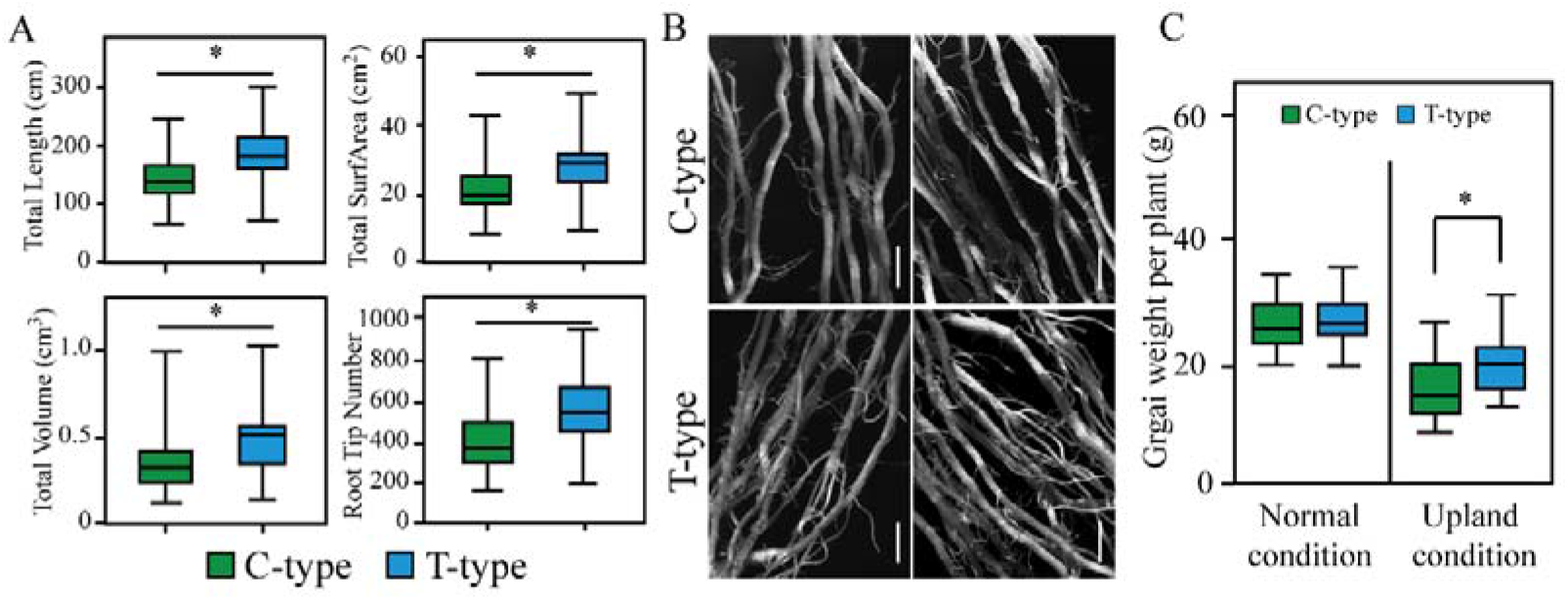
The SNP mutation (C to T) results in the functional differentiation of *OsNCED2.* A The root phenotype data, including total length, total superficial area (surface area), total root volume and root tip number, indicate that T-type plants have denser roots than C-type plants. B Comparison of the root phenotypes of C-type and T-type plants. C Grain weight per plant obtained for C-type and T-type Nipponbare plants grown under normal and upland conditions. The values in the histograms represent the means ± SDs (n = 24). * indicates significant differences between C-type and T-type plants at *p* < 0.05, as determined by Student’s *t*-test. Scale bars, 0.5 cm.

### Pathways and Downstream Genes Involved in Aerobic Adaptation Mediated by *OsNCED2*

To gain insights into the genes and pathways that contribute to aerobic adaptation and are regulated by *OsNCED2*, we screened the differentially expressed genes (DEGs) in the *OsNCED2^T^-*OE3 lines compared with the WT plants by transcriptome sequencing. As shown in Supplemental Table S4, 200 and 546 genes were up- and downregulated in the leaves of the *OsNCED2^T^*-OE3 plants compared with those of the WT plants under normal growth conditions, respectively, and 414 and 297 genes were up- and downregulated in the *OsNCED2^T^*-OE3 roots compared with the WT roots under normal growth conditions (Supplemental Table S5). Interestingly, a large number of genes encoding retrotransposon or transposon proteins were highly upregulated in both the leaves and roots of the *OsNCED2^T^*-OE3 lines (Supplemental Table S4; S5), which might influence various effects on the genome.

A GO analysis revealed that the DEGs in both the leaves and roots were functionally related to all types of pathways (Supplemental Table S6; S7). In addition, the DEGs were highly differentially enriched between the leaves and roots, which implies that the ABA molecules that are synthesized by *OsNCED2* play diverse roles in different tissues. For example, many DEGs in the leaves were specifically enriched in “transcription factor activity”, “calcium ion binding”, “protein ubiquitination”, “oxidoreductase activity” and “oxidoreductase activity” etc, whereas some DEGs in the roots were enriched in “response to stress”, “nucleoside-triphosphatase activity” and “actin cytoskeleton” etc (Supplemental Table S6; S7). A further KEGG analysis also demonstrated that the DEGs identified in the *OsNCED2^T^*-OE3 were involved in “Brassinosteroid biosynthesis” “MAPK signalling pathway” and “Starch and sucrose metabolism” and in “Diterpenoid and Flavonoid biosynthesis”, respectively (Supplemental Table S8), which indicates that ABA inhibits leaf growth and development but enhances the development of roots and their response to environment stress. Particularly, a protein phosphatase 2C (PP2C, LOC_Os09g15670) encoding gene in MAPK signalling pathway is down-regulated in the leaves of *OsNCED2^T^*-OE3 lines (Supplemental Fig. S6), which relieve the inhibition to the SnRK2s and activate ABF transcript factors to close stomata (Lin et al., 2020; Park et al., 2009).

## DISCUSSION

Plants modify their development to adapt to the environment and thus protect themselves from detrimental conditions by triggering a variety of signalling pathways. The hormone ABA is usually associated with major plant responses to stress, and its biosynthesis and signalling pathways have been well characterized (Chen et al., 2019). 9-Cis-epoxycarotenoid dioxygenase encoded by the *NCED* gene oxidatively cleaves both 9’-cis-neoxanthin and 9′-cis-violaxanthin to xanthoxin, which plays vital roles in ABA biosynthesis (Anstis et al., 1975; Schwartz et al., 1997). In this study, the phylogenetic tree of both rice *NCED* paralogous and orthologous genes indicated that rice *NCED* originated from two ancestral genes that might produce orthologous genes via gene duplication stemming from a proto-*NCED* gene. *OsNCED2* is a 9-cis-epoxycarotenoid dioxygenase-encoding gene identified that exhibits high differentiation between irrigated rice and upland rice. In our previous study, we found that the SNP-T allele (*OsNCED2^T^*) is fixed in upland rice and associated with a higher ABA content and a denser root system in both natural and segregating populations (Lyu et al., 2013). Consistently, the results from this study further confirmed the function of *OsNCED2^T^* in ABA biosynthesis and root development as well as its downstream responses to aerobic environment stress.

The SNP site evolved from a C residue in irrigated rice to a T residue in upland rice under upland selection and contributes to plant aerobic adaptation. Although sequence differences, such as SNPs and indels, were detected between the upland rice and irrigated rice varieties, the expression of *OsNCED2* was not significantly different between the two ecotypes under both normal and drought conditions, which implies that the amino acid change in the SNP site might have resulted in functional variation. Both OsNCED2^T^ and OsNCED2^C^ have functions in catalysing ABA synthesis, and OsNCED2^T^ might exhibit higher enzyme activity and thus induce elevated ABA levels in both leaves and roots of upland rice. Additional studies are needed to determine whether the expression of *OsNCED2^T^* exhibits tissue specificity and plays cell type-dependent roles.

Under aerobic conditions with drought stress, defects in osmotic homeostasis and the ion balance of plant cells trigger a signal transduction pathway for the recovery of cellular homeostasis and the repair of damaged proteins and membrane systems (Viswanathan et al., 2004). In plants, MAPKs play important roles in signal transduction in response to drought, reactive oxygen species (ROS), pathogen defence, wounding, and low temperature (Jonak et al., 1996; Mizoguchi et al., 1997). Particularly a down-regulated PP2C gene related to the MAPK signalling pathway, which indicates that ABA-dependent downstream pathways in *OsNCED2^T^*-OE3 plants such as AREB/ABF transcription factors might be regulated by active SnRK2s, result in rapid stomatal closure (Lin et al., 2020; Park et al., 2009).

As part of the adaptive response to stress, ABA can inhibit vegetative growth and thus oppose the growth-promoting properties of BRs (Clouse, 2016; Wang et al., 2018). BR-related genes were markedly downregulated in *OsNCED2^T^*-OE3 plants, which suggests that *OsNCED2* might also be involved in the inhibition of plant growth and development through antagonization of the BR pathway.

To improve aerobic adaption with drought resistance, plants develop deep and denser roots and undergo rapid stomatal closure to absorb water from soil, which have positive effects on drought avoidance (Fukai and Cooper, 1995; Wang et al., 2020b; Yoshida and Hasegawa, 1982; Zhou et al., 2020). Our results indicate that the aerobic adaptation of upland rice involves an enhanced root system due to *OsNCED2^T^-*mediated improvements in root development and rapid stomatal closure, which partially endow the *OsNCED2^T^* containing rice with increased drought avoidance.

The aerobic with DT system, including osmotic adjustment, membrane-protection proteins and redox homeostasis maintenance, also plays vital roles in plant survival and reproduction. DEGs involved in osmotic adjustment-related functions, such as carbohydrate metabolism and ion transport, were also enriched in the *OsNCED2^T^*-OE3 plants. In plants, abiotic stress can induce the production of cellular ROS, which have dual functions dependent on the amount: at a low level, ROS trigger defences and developmental responses at early stages, and at a high level, ROS attack the cell membrane to break down the defence barrier and thus destroy the cell (Guo et al., 2018; Jaspers and Jaakko, 2010; Miao et al., 2006). The *OsNCED2^T^-OE3* plants strengthened their antioxidant systems, including enzymatic and nonenzymatic antioxidants, such as superoxide dismutase, catalase, ascorbate peroxidase, glutathione reductase, ascorbic acid, tocopherol, glutathione, and phenolic compounds, which can scavenge or reduce excessive deleterious ROS, in response to an imbalance between ROS production and antioxidant defence (Gupta et al., 1993; Sharma and Dietz, 2009), which indicates that *OsNCED2^T^* contributes to DT through the maintenance of redox homeostasis in plants.

Unlike mobile animals, sessile plants cannot evade but rather have to tolerate unfavourable environmental conditions, such as drought, salinity and cold. Our results reveal a key gene that functions in the ABA biosynthesis pathway and subsequent ABA signalling pathway and contributes to ABA-triggered DA and DT. Consistent with the viewpoint that ABA increases resistance to abiotic stresses in land plant evolution and terrestrial adaptation, which originated or expanded in the common ancestor of Zygnematophyceae and embryophytes (Cheng et al., 2019; Wang et al., 2020a; Zhang et al., 2020), it is conceivable that the ABA responses meditated by *OsNCED2^T^* might offer an advantageous strategy used by upland rice to survive and breed under aerobic conditions. Furthermore, *OsNCED2^T^*-containing rice exhibit a higher yield under both rainfed upland and severe drought conditions, and these findings provide new genetic basis for the breeding of aerobic adaptation rice.

## MATERIALS AND METHODS

### Plant Materials, Growth Conditions and Stress Treatments

*Oryza sativa* L. cv. Nipponbare and Koshihikari were used in this study as the WT and background rice, respectively. Three NILs containing the *OsNCED2^T^* locus (named T-Koshihikari-1~3) were obtained by crossing the upland variety IRAT104 with the irrigated variety Koshihikari and selected from the BC_4_F_9_ population lines with Koshihikari as the recurrent parent by MAS with a CAPs marker (Supplemental Table S2).

For phenotype analysis at the seedling stage, seeds were grown in Hoagland’s nutrient solution under controlled conditions in a growth chamber with 14-h of daylight at 28 °C and a 10-h dark period at 25 °C. PEG6000 was added to the nutrient solution to a final concentration of 20% to simulate drought stress. The transgenic rice and *OsNCED2^T^*-NIL plants and their background parents, Nipponbare and Koshihikari, were evaluated in term of their root system and DT performances. Phenotyping was also performed in the field under both irrigated and upland conditions in Xishuangbanna, Yunnan Province. For irrigated condition, seeds were germinated in a seedbed, and the resulting seedlings were transplanted to a paddy field, where water was ponded on the soil surface throughout the growth and developmental period. For upland condition, we conducted direct seeding by dibbling the seeds in dry soil, during growth stages, no any irrigation was applied but depend on rainfall. Another aerobic with drought stress treatment experiment was performed using waterproof installations at the breeding centre of Yunnan University. For each line, we planted three replicates, and each replicate consisted of 12 individuals in two rows (six individuals in each row), with a row spacing of 20 centimetres and a plant spacing of 20 centimetres. From each line, approximately eight individuals were randomly selected and phenotyped. Leaf tissues were harvested at the indicated times and stored at −80 °C for further analysis.

### Measurement of ABA

For the ABA content assays, the shoots (containing the coleoptile and first leaves) and roots of seedlings were harvested. For each sample, approximately 0.2 g of fresh tissue was homogenized under liquid nitrogen, weighed, and extracted for 24 h with cold methanol containing antioxidant and 6 ng of ^2^H_6_-ABA (internal standard; OlChemIm). The endogenous ABA was purified and measured as previously described (Fu et al., 2012) with some changes in the detection conditions (Yin et al., 2015).

### RNA Extraction and Quantitative Real-time PCR (qRT-PCR)

Total RNA from rice tissues of the three genotypes was extracted using the TRIzol reagent (Invitrogen, USA). cDNA synthesis was performed using EasyScript First-strand cDNA Synthesis SuperMix (TransGene, Beijing, China) according to the manufacturer’s instructions. qRT-PCR analysis was carried out using TaKaRa SYBR Premix Ex Taq^TM^ according to the manufacturer’s instructions. The relative expression of each gene was calculated using the 2^−ΔCt^ and 2^−ΔΔCt^ method (Livak and Schmittgen, 2001). The primers (Supplemental Table S2) used for qRT-PCR were subsequently tested using a dissociation curve analysis, and the results confirmed the absence of nonspecific amplification. The ubiquitin gene was used as the reference. All the analyses were performed with three biological replicates.

### Vector Construction and Genetic Transformation

The coding region of *OsNCED2* was amplified from the genomic DNA of two types of rice (IRAT104 and Nipponbare) by PCR using *Pst* I and *Spe* I linker primers (Supplemental Table S2). The resulting *OsNCED2* fragments were inserted into the *Pst* I and *Spe* I sites of pCUbi1390 (Peng et al., 2009) to generate *UbiPro::OsNCED2^T^* and *UbiPro::OsNCED2^C^*, respectively. Fragments of 3.4 kb in the *OsNCED2* promoter of IRAT104 were amplified from rice genomic DNA (Nipponbare) by PCR and inserted into the pMDC162 vector (Curtis and Grossniklaus, 2003) using Gateway technology to generate *OsNCED2Pro::GUS*. All the vectors were introduced into *Agrobacterium tumefaciens* strain *EHA105* and then transferred into Nipponbare plants via *Agrobacterium*-mediated transformation in accordance to the standard protocol (Duan et al., 2012).

### Physiological Traits of the Three Genotypes under Drought Stress

Fresh leaves (0.5 g) of the seedlings under normal and stress conditions were harvested and used to measure the MDA content and the SOD, POD and CAT activity levels according to described methods (Shin et al., 2012; Shukla et al., 2012). The RWC was calculated using the following formula: RWC (%) = [(FM − DM)/(TM − DM)] × 100, where FM, DM, and TM are the fresh, dry, and turgid masses of the weighed leaves, respectively. The relative electrolyte leakage (REL) or solute leakage from the sampled rice leaves was evaluated using the method described by Arora et al. (Arora et al., 1998). The percent injury induced by each treatment was calculated from conductivity data using the following equation: % injury = [(% L(t)-% L(c))/(100-% L(c))] × 100), where % L (t) and % L(c) are the percent conductivity of the treated and control samples, respectively. The second leaves from 1-hour drought-stressed plants were used for SEM analyses, as described by (You et al., 2012) with minor modifications. Fresh leaf samples were prefixed for 3 h in 3% glutaraldehyde-sodium phosphate buffer (0.1 M) at room temperature and rinsed three times with 0.1 M sodium phosphate buffer. Postfixation was performed with 2% OsO_4_ at 4°C. The samples were dehydrated through an ethanol series and infiltrated with an isoamyl acetate series. The samples were then coated with metal particles for SEM analysis to observe the guard cells. A Hitachi S750 scanning electron microscope (http://www.hitachi-hitec.com/global/em/) was used to obtain photographs, and the numbers of guard cells in randomly selected fields were counted and statistically analysed.

### Subcellular Localization of GFP-OsNCED2 Fusion Proteins

The open reading frames (ORFs) of *OsNCED2* were inserted into pMDC43 as C-terminal fusions to the green fluorescent protein (GFP) reporter gene driven by the CaMV 35S promoter (Curtis and Grossniklaus, 2003). For transient expression, the *GFP-OsNCED2* fusion vector constructs were transformed into the leaves of 3-week-old tobacco (*Nicotiana benthamiana*) by *A. tumefaciens* infiltration (Sparkes et al., 2006). The green fluorescence resulting from GFP-OsNCED2 expression was observed using a confocal laser scanning microscope (LSM700, Zeiss, Jena, Germany). The 35S::GFP construct was used as a control.

### Histochemical GUS Assay

For GUS staining analysis, sample tissues were submerged in GUS staining buffer (containing 2 mM 5-bromo-4-chloro-3-indolyl glucuronide, 0.1 M sodium phosphate buffer [pH 7.0], 0.1% [v/v] Triton X-100, 1 mM potassium ferricyanide, 1 mM potassium ferrocyanide, and 10 mM EDTA), vacuum infiltrated for 10 min, and then incubated overnight at 37 °C. The staining buffer was removed, and the samples were then cleared with 95% (v/v) ethanol and observed using a stereoscope (LEICA, 10447157, Germany).

### Gene Editing for the SNP Mutation Site in *OsNCED2*

The 23-bp targeting sequences (including PAM) (Supplemental Table S2) were selected within the target gene *OsNCED2*, and their targeting specificity was confirmed through a BLAST search against the rice genome (http://blast.ncbi.nlm.nih.gov/Blast.cgi). The designed targeting sequences were synthesized and annealed to form oligo adaptors. The pCSGAPO1 vector (Cong et al., 2013) was digested with *Bsa* I and purified using a DNA purification kit (Tiangen, China). A ligation reaction (10 μl) containing 10 ng of the digested pCSGAPO1 vector and 0.05 mM oligo adaptor was conducted and directly transformed into *E. coli* competent cells to produce CRISPR/Cas9 plasmids. The CRISPR/Cas9 plasmids were introduced into A. *tumefaciens* strain EHA105. The transformation of rice and the genotyping of the SNP site of *OsNCED2* were performed as described above.

### RNA-Seq Analysis

Total RNA from rice seedlings belonging to the *OsNCED2^T^*-OE3 and WT lines was extracted using the TRIzol reagent according to the manufacturer’s instructions (Invitrogen, Waltham, MA, USA). For each sequencing library, 100 mg of RNA from each replicate was mixed together. The libraries were sequenced using an Illumina HiSeq 2000 Sequencing System. Low-quality nucleotides (< Q20) were trimmed from the raw sequences obtained for each sample, and pair-end reads with at least one end with a length < 30 bp were removed using an in-house Perl script. The retained high-quality reads were mapped to the Michigan State University Rice Genome Annotation Project database (ftp://ftp.plantbiology.msu.edu/pub/data/Eukaryotic_Projects/o_sativa/annotation_dbs/pseudomolecules/version_7.0) using Bowtie (Ouyang et al., 2007). The Cuffdiff module was used for identification of the DEGs. Chi-square tests were used to identify genes with significant differences in their relative abundance levels (as reflected by the total counts of individual sequence reads) between two samples using IDEG6 software based on the criteria p ≤ 0.001 and fold change > 2 (Romualdi et al., 2003; Z.N. et al., 2003). The pathway and GO enrichment analyses of rice DEGs were conducted using EXPath 2.0 (Chien et al.; Jia et al., 2018).

## Supplemental Data

**Supplemental Figure S1.**
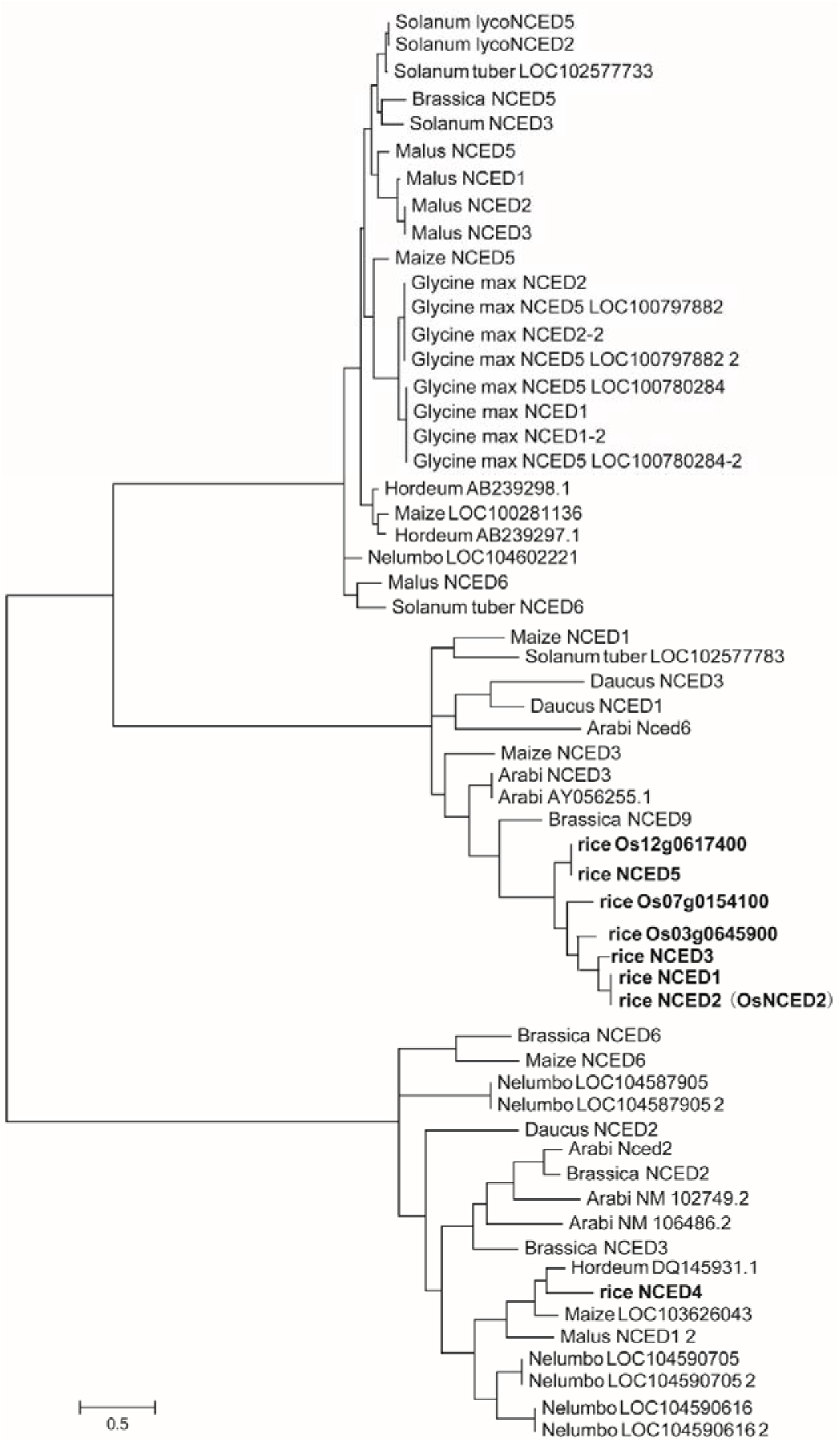
Phylogenetic analysis of the NCED proteins in plants. The maximum parsimony tree of the NCED proteins from various plants was based on the sequence database in the UniProt Knowledgebase (UniProtKB).

**Supplemental Figure S2.**
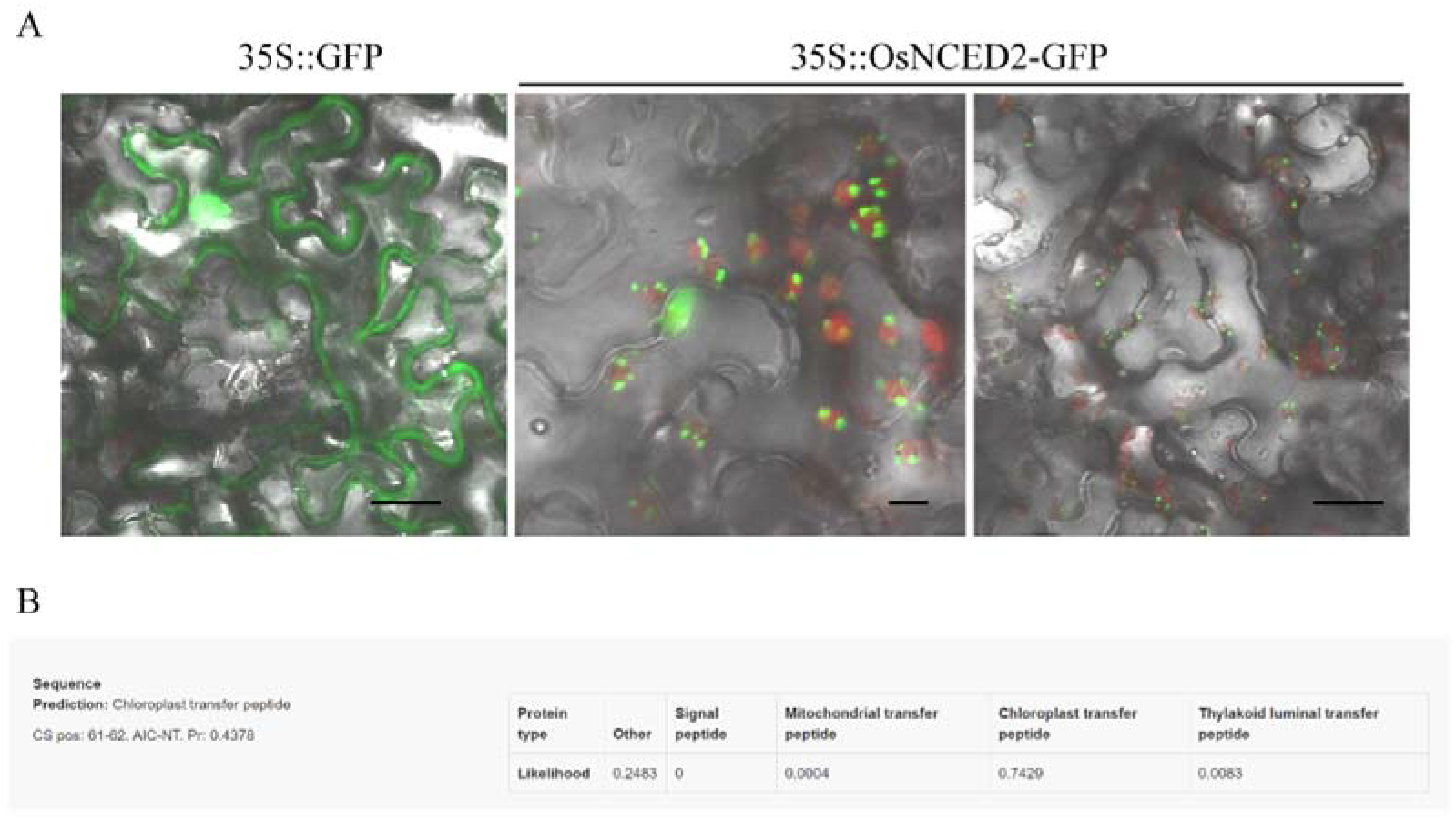
Subcellular localization analysis of OsNCED2 in plants. A GFP was extensively localized in both the nucleus and cytoplasm in *N. benthamiana* protoplasts, while OsNCED2-GFP was positively localized in the chloroplast. The chlorophyll fluorescence is shown in red. The scale bars represent 10 μm. B Prediction of subcellular location via Target P analysis.

**Supplemental Figure S3.**
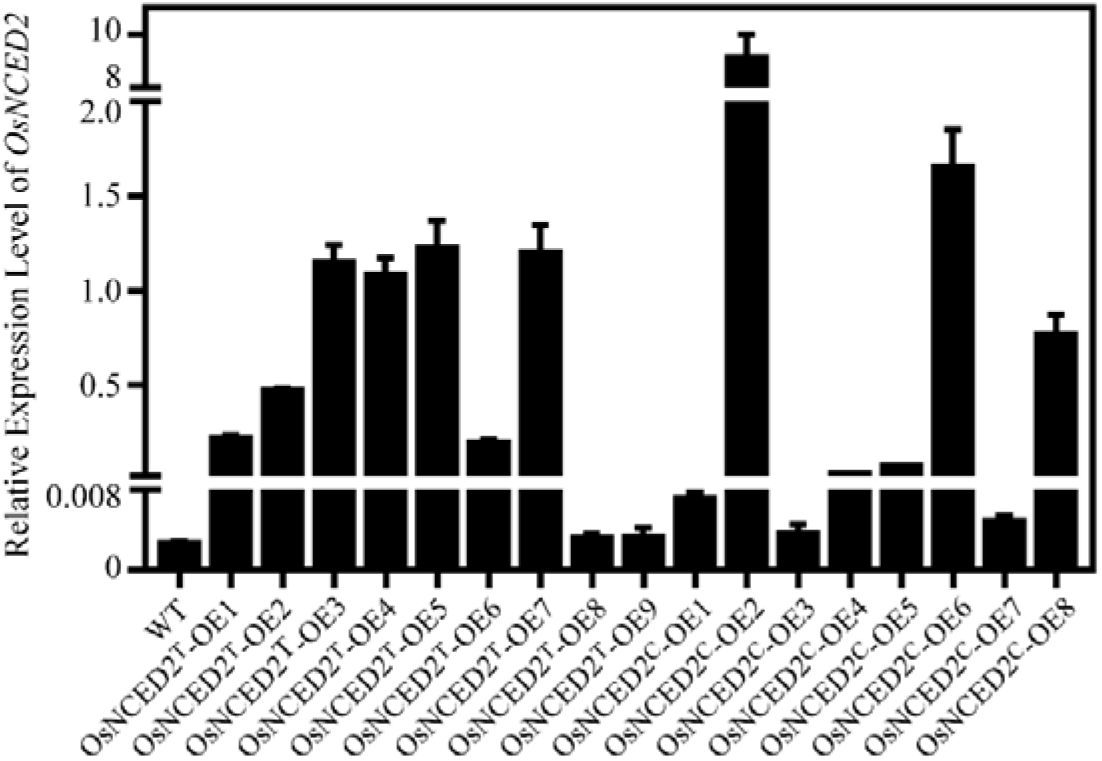
Analysis of *OsNCED2* expression in transgenic plants overexpressing *OsNCED2^T^* and *OsNCED2^C^*.

**Supplemental Figure S4.**
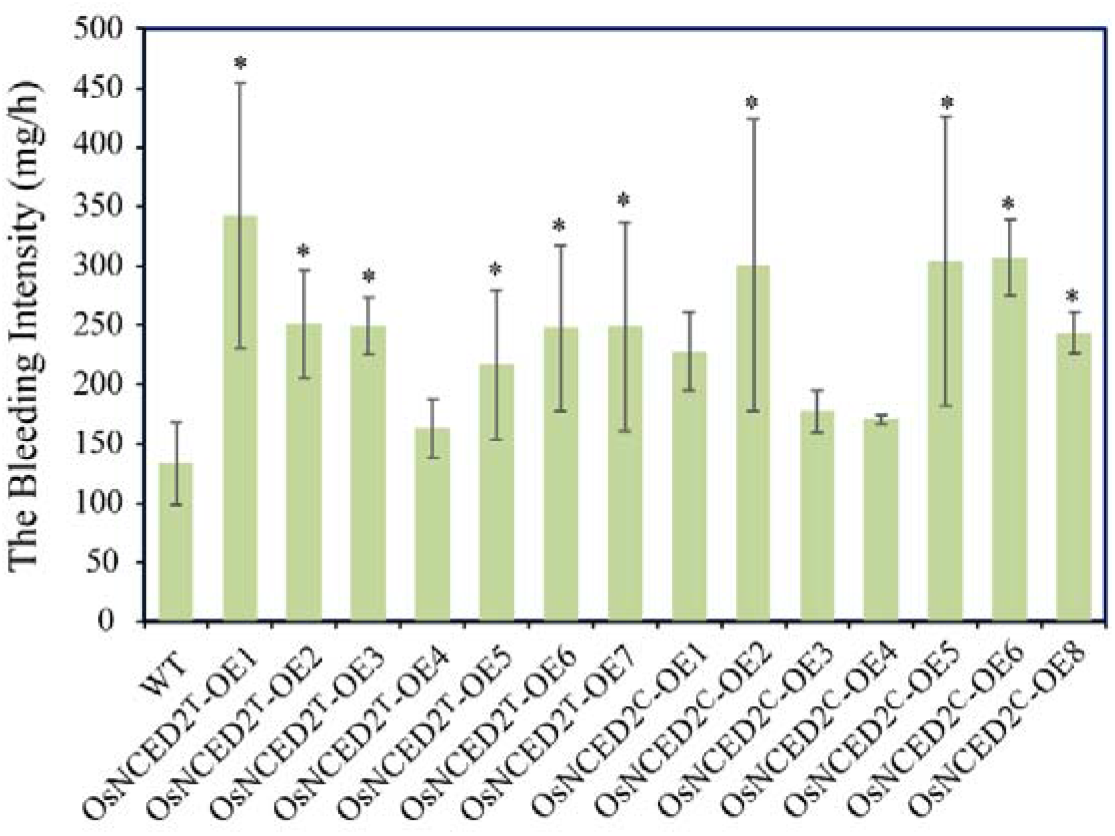
Bleeding intensity of transgenic plants overexpressing *OsNCED2^T^* and *OsNCED2^C^*.

**Supplemental Figure S5.**
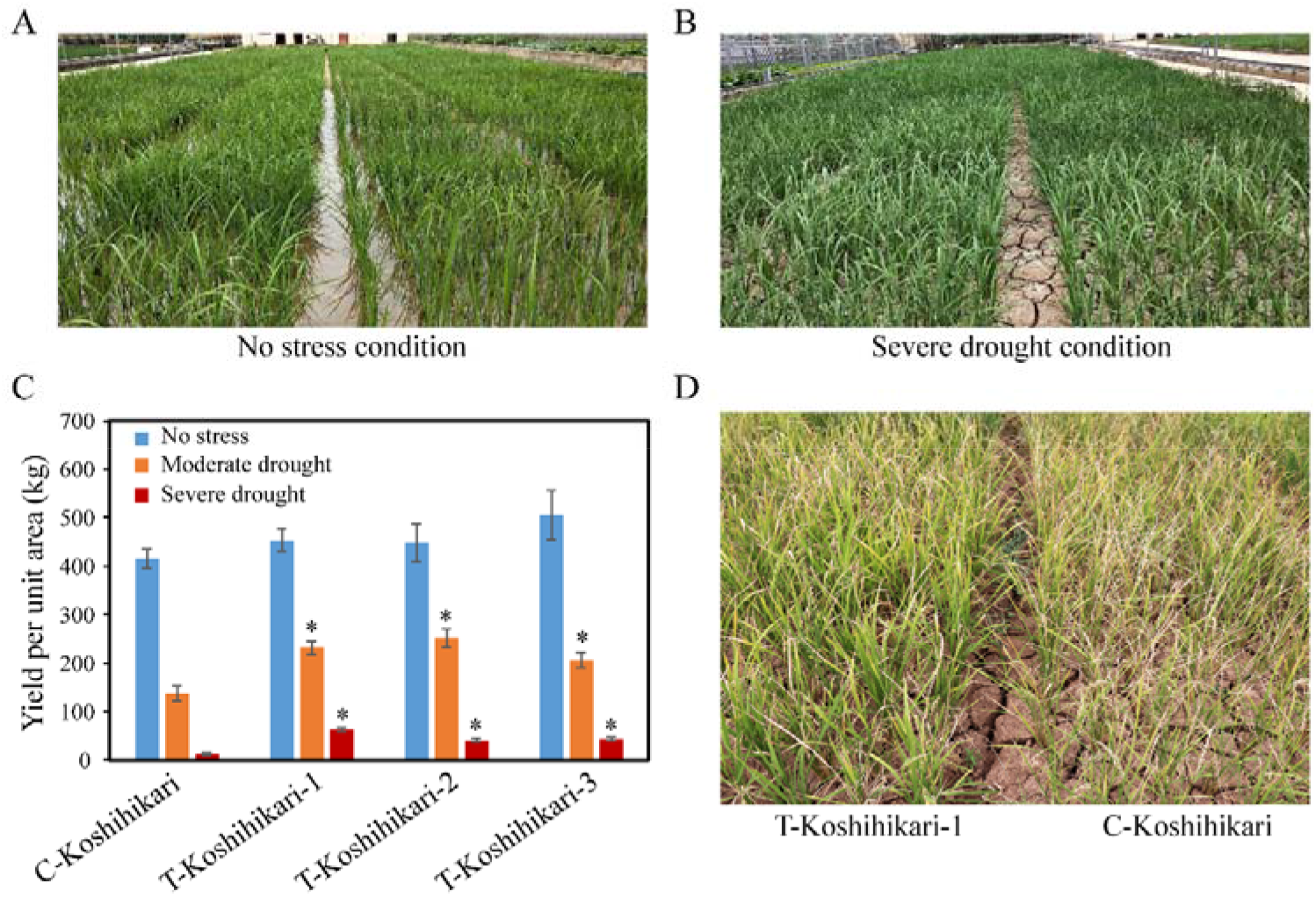
Phenotyping of *OsNCED2^T^*-NIL and C-Koshihikari plants in both paddy fields and aerobic conditions (severe-drought conditions under a rainout shelter). Comparisons of rice performance in paddy fields (A) and aerobic land (B). C Yield per unit area of *OsNCED2^T^*-NIL and C-Koshihikari plants. * indicates a significant difference from the C-Koshihikari plants at *p* < 0.05, as determined by Student’s *t*-test. D Performance of *OsNCED2^T^*-NIL and C-Koshihikari plants in drylands at the maturation stage.

**Supplemental Figure S6.**
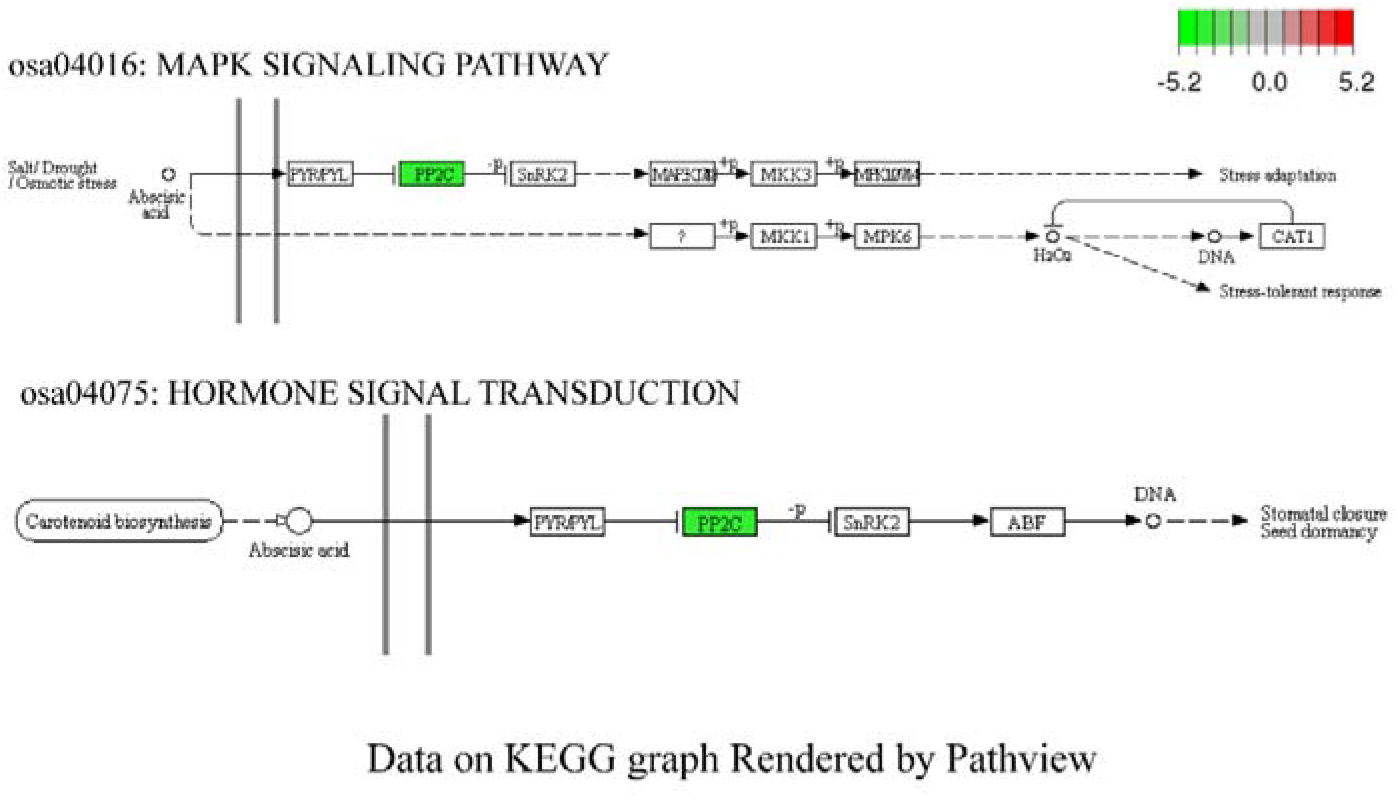
MAPK signaling pathway and hormone signal transduction pathway rendered by Pathview.

## Supplemental Tables

**Supplemental Table S1.** *Cis*-acting elements in the *OsNCED2* promoter.

**Supplemental Table S2.** Information on the primers used in this study.

**Supplemental Table S3.** Agronomic traits of *OsNCED2^T^*-NIL plants.

**Supplemental Table S4.** DEGs in the leaves of *OsNCED2^T^*-OE3, when compared to WT under the normal growth conditions.

**Supplemental Table S5.** DEGs in the roots of *OsNCED2^T^*-OE3, when compared to WT under the normal growth conditions.

**Supplemental Table S6.** Function categories list of leaves DEGs in *OsNCED2^T^*-OE3 compared with WT via GO analysis.

**Supplemental Table S7.** Function categories list of roots DEGs in *OsNCED2^T^*-OE3 compared with WT via GO analysis.

**Supplemental Table S8.** List of pathways and related DEGs enriched in *OsNCED2^T^*-OE3 vs WT via KEGG analysis.

## ACKNOWLEDGEMENTS AND FUNDING

This work was supported bygrants fromthe National Natural Science Foundation of China (U1602266 and31601274) and grants from the Yunnan Provincial Science and Technology Department (2018IC096, 2019ZG013 and 2019HC028).

## AUTHOR CONTRIBUTIONS

F.Y.H. and L.Y.H. conceived the project. L.Y.H. designed and performed the experiments. Y.C.B., S.W.Q. and M.N. performed the molecular biologyexperiments, including vector construction and the transformation and genotyping of transgenic rice. S.L.Z., G.F.H. and J.Z. performed the phenotyping analysis. W.S.W. and B.Y.F. performed the RNA-Seq data analysis. L.Y.H. and F.Y.H. wrote the manuscript, and J.L. revised the draft. All the authors read and approved the final manuscript.

